# A comprehensive portrait of cilia and ciliopathies from a CRISPR-based screen for Hedgehog signaling

**DOI:** 10.1101/156059

**Authors:** David K. Breslow, Sascha Hoogendoorn, Adam R. Kopp, David W. Morgens, Brandon K. Vu, Kyuho Han, Amy Li, Gaelen T. Hess, Michael C. Bassik, James K. Chen, Maxence V. Nachury

**Author notes:** Co-first author (David K. Breslow, Sascha Hoogendoorn). Correspondence David K. Breslow, James K. Chen, Maxence V. Nachury.

## Abstract

The primary cilium organizes Hedgehog signaling, shapes embryonic development and is the unifying cause of the ciliopathies. We conducted a functional genomic screen for Hedgehog signaling by engineering antibiotic-based selection of Hedgehog-responsive cells and applying genome-wide CRISPR-mediated gene disruption. The screen robustly identifies factors required for ciliary signaling with few false positives or false negatives. Characterization of hit genes uncovers novel components of several ciliary structures including a protein complex containing ε- and δ- tubulin that is required for centriole maintenance. The screen also provides an unbiased tool for classifying ciliopathies and reveals that many forms of congenital heart defects are ciliopathies. Collectively, this screen enables a systematic analysis of ciliary function and of ciliopathies and also defines a versatile platform for dissecting signaling pathways through CRISPR-based screening.

## Introduction

The primary cilium is a surface-exposed microtubule-based compartment that serves as an organizing center for diverse signaling pathways^1-3^. Consistent with a central role for cilia in signaling, mutations affecting cilia cause ciliopathies, a group of developmental disorders that includes Joubert Syndrome, Meckel Syndrome (MKS), Nephronophthisis (NPHP), and Bardet-Biedl Syndrome (BBS). The defining symptoms of ciliopathies include skeletal malformations (e.g. polydactyly), mental retardation, sensory defects (e.g. retinal degeneration), obesity, and kidney cysts and are thought to arise from misregulation of ciliary signaling pathways. Over the past fifteen years, advances in human genetics have led to the identification of over 70 ciliopathy disease genes^4^. However, the molecular basis for many ciliopathy cases remains undiagnosed, suggesting that additional ciliopathy genes have yet to be identified^5^. More broadly, many mechanistic aspects of cilium assembly and function remain poorly understood, indicating a need for systematic discovery and characterization of genes that enable ciliary signaling.

A key paradigm for ciliary signaling is the vertebrate Hedgehog (Hh) pathway, which exhibits a strict dependence on primary cilia for signaling output^3^. Remarkably, all core components of the Hh signaling machinery – from the receptor PTCH1 to the GLI protein transcriptional effectors – dynamically localize to cilia during signal transduction. Nonetheless, the precise role of cilia in Hh signaling remains elusive, and each successive step in signal transduction is still incompletely understood^6^. Given the key roles of Hh signaling in embryonic development and in cancers such as medulloblastoma and basal cell carcinoma^7,8^, there is a pressing need to fill the gaps in our understanding of Hh signal transduction.

Efforts to systematically identify genes needed for cilium assembly or Hh signaling have been reported by a number of groups. However, these studies have relied on arrayed siRNA libraries and exhibit the high rates of false positives and false negatives that are characteristic of RNA interference (RNAi)-based screens^9-12^. Recently, genome-wide screening using the CRISPR/Cas9 system for gene disruption has emerged as a powerful tool for functional genomics^13-16^. CRISPR-based screens have already aided greatly in defining genes required for proliferation of mammalian cells, revealing a core set of essential genes and cancer cell line-specific vulnerabilities. However, the pooled screening format used in these studies requires a means to select for/against or otherwise isolate cells exhibiting the desired phenotype, a technical requirement that has significantly limited the scope of biological applications amenable to pooled CRISPR-based screens. Indeed, most studies to date have searched for genes that either intrinsically affect cell growth or that affect sensitivity to applied toxins, drugs, or microorganisms^17-24^.

Here, we engineered a Hh pathway-sensitive reporter to enable the systematic identification of genes that participate in Hh signal transduction and primary cilium function. Combining this reporter with a single guide RNA (sgRNA) library targeting the mouse genome, we conducted a CRISPR-based screen that systematically identified ciliary components, Hh signaling machinery, and ciliopathy genes with few false positives or false negatives. Our screen also revealed many genes that had not previously been characterized or linked to ciliary signaling, and we show here that these hits include new components of cilia and centrioles and novel ciliopathy genes.

## Results

### Development of a Hh pathway transcriptional reporter for pooled screening

Pooled functional screening requires the ability to enrich or deplete mutant cells that exhibit a desired phenotype from within a large population of cells. Because ciliary signaling is not intrinsically linked to such a selectable/isolatable phenotype, we engineered a reporter construct that converts Hh signaling into antibiotic resistance (Fig. 1A-B). This transcriptional reporter was introduced into mouse NIH-3T3 fibroblasts, a widely used model cell line for Hh signaling and cilium biology^25^. These cells were then further modified to express the Cas9 endonuclease fused to BFP (3T3-[Shh-BlastR;Cas9] cells).

To confirm that our reporter cell line faithfully recapitulates ciliary Hh signaling and to test its suitability for CRISPR-based mutagenesis, we introduced sgRNAs targeting regulators of the canonical Hh pathway (Supplementary Table 1). *Smo*, a key Hh pathway transducer, and *Ift88*, an intraflagellar transport (IFT) complex subunit needed for cilium assembly, are required for Hh signaling while GLI3 repressor dampens the Hh response and SUFU prevents Hh signaling in the absence of ligand (Fig. 1c, left). As expected, sgRNAs targeting *Smo* or *Ift88* severely reduced Sonic Hedgehog N-terminal domain (ShhN)-induced blasticidin resistance, deleting *Gli3* potentiated blasticidin resistance in response to ShhN, and targeting *Sufu* gave rise to ligand-independent blasticidin resistance (Fig. 1c, right). The effects of these sgRNAs on blasticidin resistance were paralleled by concordant changes in endogenous pathway outputs, including the induction of GLI1 and the shift from production of GLI3 repressor to GLI3 activator that normally occur upon pathway activation (Supplementary Fig. 1a).

**Figure 1.**
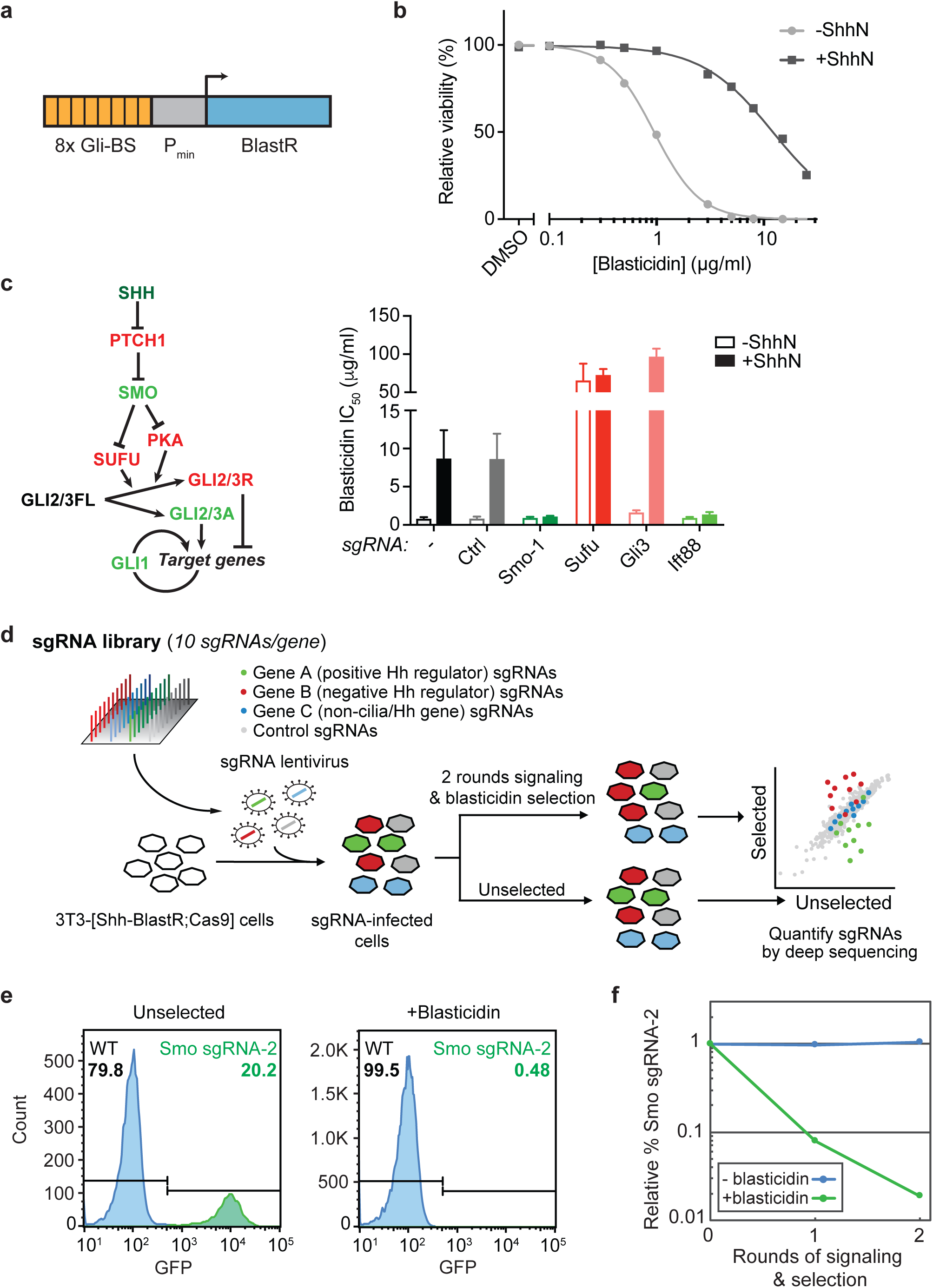
Development of a Hedgehog pathway reporter-based screening strategy. **a**) A transcriptional reporter attaches 8 copies of the GLI binding sequence (Gli-BS) to a minimal promoter (P_min_) to convert Hh signals into blasticidin resistance. **b)** Blasticidin resistance was assayed across a range of concentrations in stimulated (+ShhN) and unstimulated 3T3-[Shh-BlastR;Cas9] cells. **c**) Overview of the Hh pathway, with key negative and positive regulators shown in red and green, respectively (left). Effects of control sgRNAs on blasticidin resistance in stimulated and unstimulated 3T3-[Shh-BlastR;Cas9] cells (right). Mean inhibitory concentration 50 (IC50) values and standard deviations are shown. N = 2 to 5 independent experiments, each performed in duplicate. **d**) Overview of the screening strategy. Cells receiving a negative control (Ctrl) sgRNA, a positive regulator-targeting sgRNA, and a negative regulator-targeting sgRNA are shaded grey, green, and red, respectively. **e**) Flow cytometry histograms of cell mixtures showing the fraction of GFP positive (Smo sgRNA-2, green) cells either in the absence of selection (left) or after two rounds of signaling and selection (right). Cells expressing *Smo*-targeting sgRNA-2 are specifically depleted in the signaling/ selection conditions. **f**) Quantification of cell depletion as in (**e**).

Having established a cell line that converts differences in ciliary Hh signaling into different levels of blasticidin resistance, we next tested its suitability for pooled screening. The analysis of our pooled screen is based on quantifying sgRNAs in blasticidin-selected and unselected cell pools to identify sgRNAs that confer a selective advantage or disadvantage (Fig. 1d). We therefore mimicked screening conditions by mixing GFP-marked cells transduced with a *Smo*-targeting sgRNA with mCherry-marked cells transduced with a portion of our genome-wide sgRNA library. Monitoring GFP^+^/*Smo* sgRNA cells by flow cytometry, we found the fraction of *Smo* sgRNA cells decreased by >12-fold and by >50-fold after one and two rounds of signaling and selection, respectively (Fig. 1e-f); such changes are readily detectable by deep sequencing.

### Genome-wide screening

We conducted our genome-wide screen using a newly developed sgRNA library targeting the mouse genome^26^. Key features of this library are the use of 10 sgRNAs per gene and the inclusion of > 10,000 negative control sgRNAs that either do not have targets in the mouse genome or that target “safe” sites with no predicted functional role. We introduced this library into 3T3- [Shh-BlastR;Cas9] cells via lentiviral transduction at low multiplicity of infection and maintained sufficient cell numbers to ensure ~ 1000X coverage of the library (due to the large number of cells required, we conducted the screen in four batches using subsets of the library; Supplementary Fig. 2a). After sgRNA transduction, cells were exposed to ShhN for 24 h to fully stimulate Hh signaling, split into separate blastidicin-selected and unselected pools and then subjected to a second cycle of signaling and selection before harvesting and sgRNA counting by deep sequencing (Fig. 1c). Genes affecting ciliary signaling were identified by comparing sequencing reads in the blastidicin-selected versus unselected cell pools at the end of the experiment, while genes affecting proliferation were identified by comparing the plasmid sgRNA library used for lentivirus production to the sgRNA library after 15 days growth in the absence of blasticidin. For statistical analysis, a maximum likelihood method termed casTLE^27^ was used to determine a *P* value for each gene from the changes in abundance of the corresponding sgRNAs. In addition, the casTLE method also estimates an effect size corresponding to the apparent strength of the phenotype caused by knockout of a given gene.

**Figure 2.**
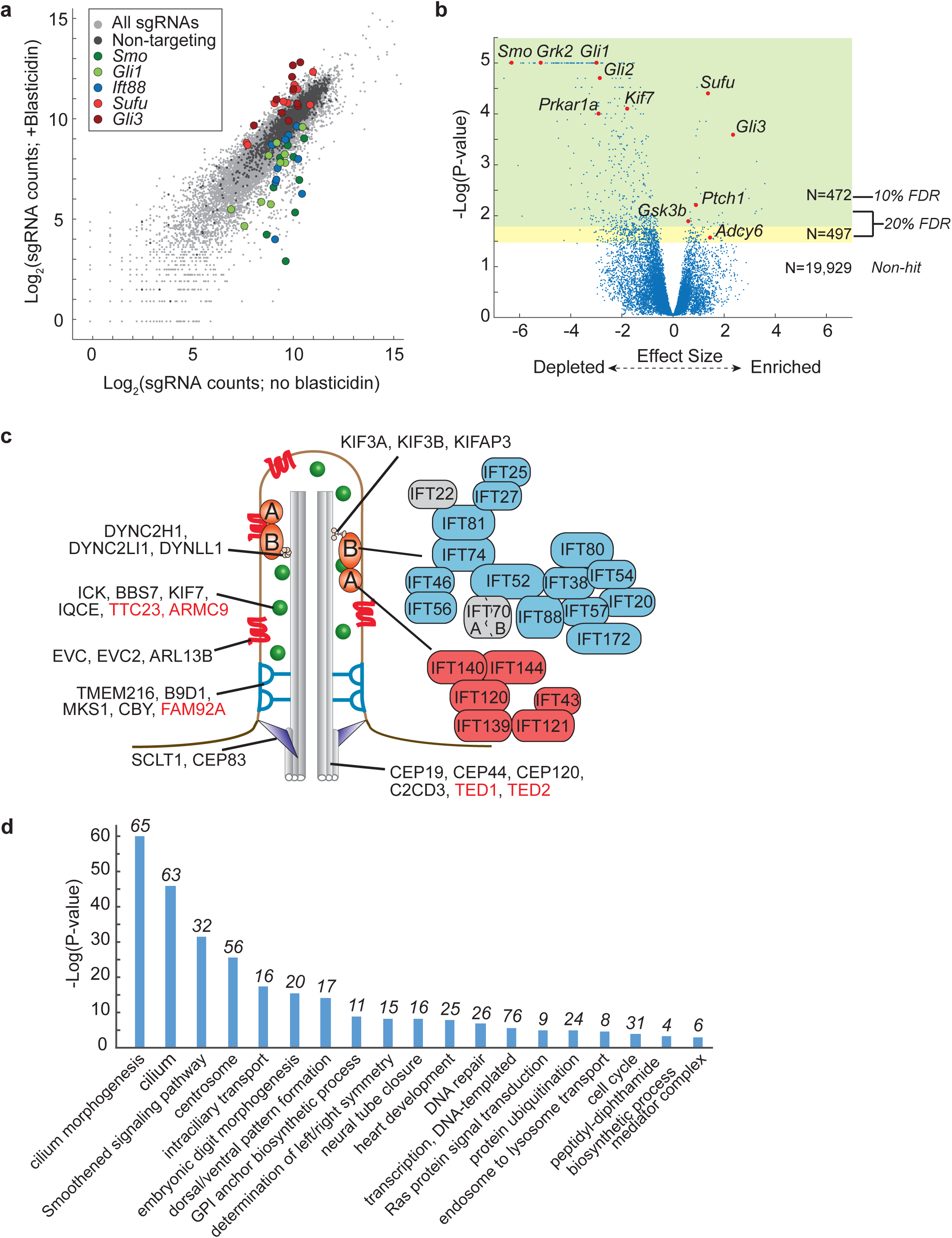
Overview of genome-wide screen results. **a**) Scatter plot showing log_2_ of normalized sgRNA counts in selected versus unselected cell pools, with sgRNAs targeting different genes shown in indicated colors. **b**) Volcano plot of casTLE *P* values against effect sizes for all genes (after filtering; see **Methods**), with core Hh pathway components in red. Green and yellow areas indicate *P*-value cutoffs corresponding to 10% and 20% false discovery rate, respectively, with the number of genes in each area indicated. **c**) Schematic illustration of a primary cilium, with known structural features and associated hit genes shown. Newly identified hit genes are in red font. For the IFT-A and IFT-B complexes, genes detected as hits are shown in red and blue boxes, respectively, while non-hit genes are in grey boxes. **d**) GO terms enriched among the top 472 hit genes. Bars indicate *P*-values from DAVID gene enrichment analysis; numbers above bars indicate number of hit genes for each class.

### Assessment of screen performance

To evaluate the quality of our screen, we first assessed its performance in detecting genes affecting growth. This readout is independent of our reporter-based selection strategy and enables comparisons to other growth-based screens. It also provides an opportunity to evaluate the performance of our sgRNA library, as this study represents its first use in a genome-wide screen. Using reference positive and negative essential gene sets^28^, we found that our screen detected growth-affecting genes with few false positives or false negatives, identifying >90% of essential genes with a 5% false discovery rate (FDR) (Supplementary Fig. 2b and Supplementary Tables 2-3). This performance is comparable to that seen recently with two other libraries^19,21^.

We next evaluated the ability of the blasticidin-based screen to identify genes known to participate in ciliary Hh signaling. Initial inspection of screen results for *Smo*, *Ift88*, *Gli1*, *Gli3*, and *Sufu* revealed several sgRNAs targeting each gene that were depleted or enriched as expected upon blasticidin selection (Fig. 2a). Virtually all known Hh signaling components were among the top hits, including positive regulators *Smo*, *Grk2*, *Kif7*, *Prkar1a*, *Gli1*, and *Gli2* and negative regulators *Ptch1*, *Adcy6*, *Gsk3b*, *Sufu*, and *Gli3* (Fig. 2b and Supplementary Table 4).

In addition to identifying Hh pathway genes, our screen recovered many hits required for the assembly and function of primary cilia. Remarkably, these hits encompass nearly all functional and structural elements of cilia, highlighting the diverse features of cilia needed for effective signaling (Fig. 2c). For example, several hits encode components of the basal body that nucleates the cilium, the transition fibers that anchor the basal body to the cell surface, the transition zone that gates protein entry into the cilium, the ciliary motors that mediate intraciliary transport, and the IFT complexes that traffic ciliary cargos (Fig. 2c and Supplementary Table 4). The detection of nearly all IFT-A and IFT-B genes as hits also underscores the low rate of false negatives.

We observed no apparent correlation between growth and signaling phenotypes, indicating that our antibiotic selection strategy is not biased by general effects on proliferation (Supplementary Fig. 2c). However, because low sgRNA counts were commonly obtained for genes with the strongest growth defects, the estimated signaling phenotypes for these genes may be less reliable and were considered separately. In total, we obtained 472 hits at a 10% FDR and 969 hits at a 20% FDR, and the majority of these hits led to decreased signaling rather than increased signaling (Fig. 2c). Gene ontology (GO) term analysis using DAVID^29^ revealed that the top 472 hit genes were significantly enriched for many expected functional categories (e.g. cilium morphogenesis, *P* < 1x10^-60^; Smoothened signaling pathway, *P* < 1x10^-31^) as well as some novel categories, indicating new avenues for investigation (Fig. 2d and Supplementary Table 5). In some cases, corroborating reports support these new connections: for example, mouse mutants for two hit genes that mediate diphthamide modification exhibit Hh pathway-related phenotypes such as polydactyly^30,31^. *DPH1* mutations have also been shown to cause a poorly characterized syndrome with several ciliopathy-like features including cerebellar vermis hypoplasia, craniofacial malformations, and cardiac defects^32^. Other unexpected gene categories, such as endosome-to-lysosome transport or the Ras signaling pathway, will require further investigation.

We next sought to use reference sets of expected hit and non-hit genes to quantitatively assess screen performance. To this end, we curated a set of ciliogenesis reference genes^9^ to generate a list of 130 expected hits (Supplementary Table 3). For expected non-hits, we used 1386 olfactory and vomeronasal receptor genes, as they are likely not expressed in cultured fibroblasts. Using these reference gene sets, we calculated precision-recall and receiver operating characteristic (ROC) curves (Fig. 3a) from the *P* values generated by casTLE. Both performance metrics show a high area under the curve (0.802 for precision-recall, 0.892 for ROC), demonstrating that our screen detects hits with high sensitivity and precision (Fig. 3a).

**Figure 3.**
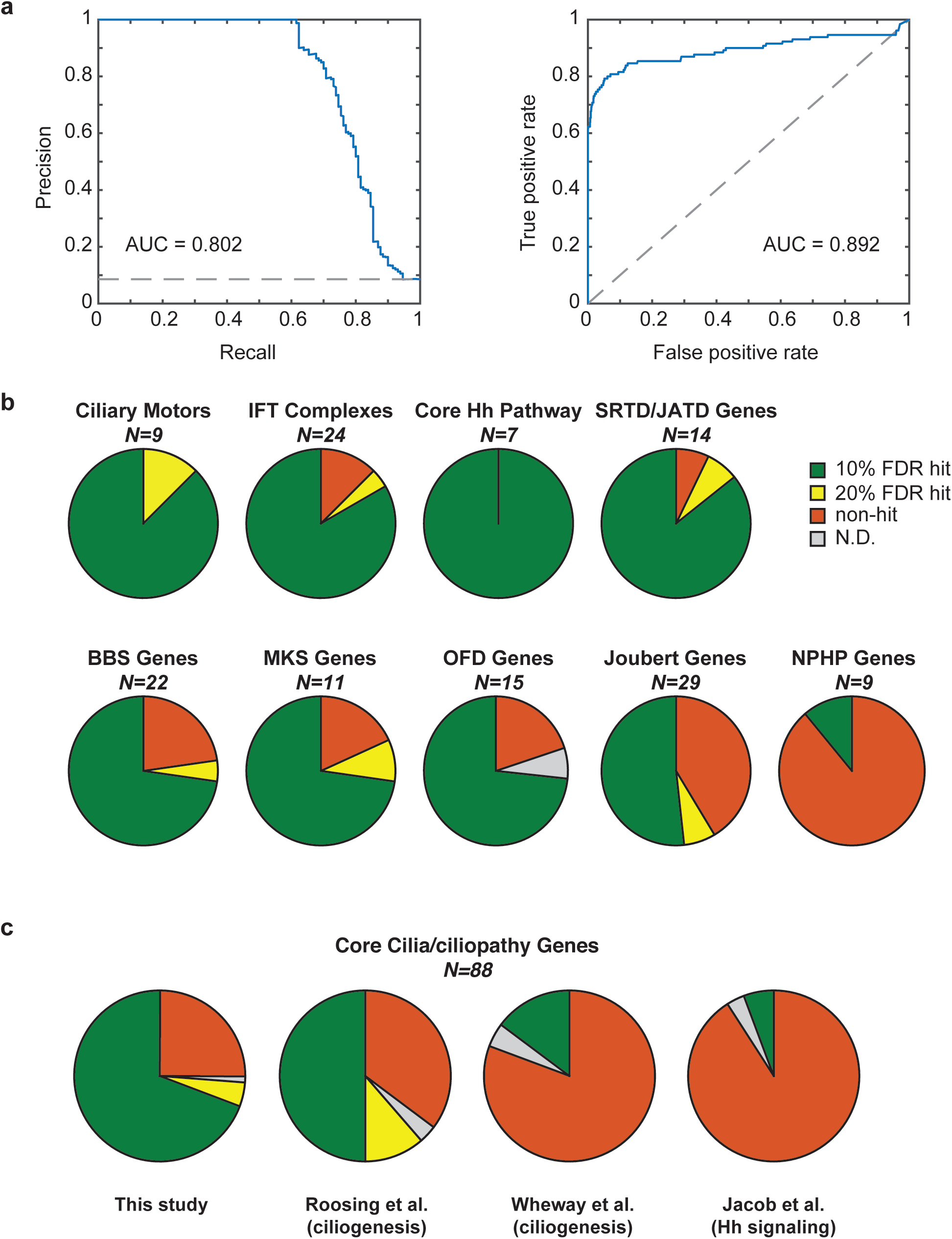
Evaluation of screen performance. **a**) Assessment of screen performance using positive and negative reference genes, as determined by precision-recall analysis (left) and ROC curve (right), with the area under each curve (AUC) shown. Dashed lines indicate performance of a random classification model. **b**) Analysis of hit gene detection for select gene categories, with the fraction of hits detected at 10% or 20% FDR, not detected, or not determined shown; see Supplementary Table 3 for details. The NPHP category includes genes mutated exclusively in NPHP and not other ciliopathies. Abbreviations: SRTD (short rib thoracic dysplasia), JATD (Jeune asphyxiating thoracic dysplasia), OFD (Oral-Facial-Digital Syndrome). **c**) Hit gene identification is compared for the indicated datasets. Pie charts show the fraction of genes detected as hits across all genes included in part (**b**), except the NPHP-specific category; see Supplementary Figure 4 for detail among individual categories.

As a second means of evaluation, we compared our ability to detect expected hit genes to that of three related screens. These studies used arrayed siRNA-based screening to study either Hh signaling using a luciferase reporter^11^ or ciliogenesis using microscopy-based measures of ciliary markers^9,10^. While there are notable differences among the screens (e.g. Roosing et al. incorporated data from other sources such as gene expression to score hits^9^), they each defined a number of hit genes similar to (or greater than) our screen, thus making it straightforward to compare performance. Overall, we detected the vast majority of expected hits across functional categories ranging from Hh pathway components to ciliopathy genes. Furthermore, even though our screen was focused on Hh signaling, we detected a greater fraction of ciliary hits than the ciliogenesis screens across categories including IFT subunits, ciliary motors, and nearly all classes of ciliopathy genes (Fig. 3b-c, Supplementary Fig. 3a and Supplementary Table 4). One exception however is the group of genes mutated in NPHP, where we found few hit genes (particularly when analyzing genes mutated exclusively in NPHP). While the basis of this finding warrants further study, it raises the possibility that NPHP pathophysiology may be distinct from that of other ciliopathies.

As a final assessment of our screening platform, we evaluated reproducibility across replicate screens. We observed high concordance among hits for the 95 genes that were measured in two different batches of the screen (Supplementary Fig. 3b), with 50 of 54 screen hits also scoring as hits in the second batch. Similarly, strong overlap in hits was found for 263 genes that were screened using two similar but distinct means to initiate Hh signaling: addition of PTCH1 ligand (ShhN) or SMO agonist (SAG) (Supplementary Fig. 3c). This reproducibility makes it possible to directly compare screens and pinpoint genes acting at specific steps in Hh signal transduction. For example, *Gas1* was a hit in the ShhN screen but not in the SAG screen, a result in agreement with GAS1’s known function as a Shh co-receptor^33,34^. Overall, our results indicate that combining CRISPR-based screening with a pathway-specific transcriptional reporter is a powerful strategy for functional genomics.

### Identification of new ciliary components

The effectiveness of our screen in identifying genes known to participate in cilium function corroborates the essentiality of this organelle for vertebrate Hh signaling, and our success in identifying the known Hh signal transduction machinery suggests that novel Hh pathway components will be found among the many hits. Given that there are likely many more cilium-related genes than core Hh pathway components, we set out to characterize the roles of six hit genes in cilium biology as a way to further establish the value of our screen.

We first focused on *Fam92a* and *Ttc23* because their gene products contain domains associated with membrane trafficking. For *Fam92a*, we confirmed our pooled screen results in cells transduced with individually cloned sgRNAs, finding that *Fam92a* knockout caused a strong defect in inducible blasticidin resistance (Supplementary Fig. 4a). This defect was also seen for induction of luciferase from a GLI binding site reporter and could be rescued by re-introduction of sgRNA-resistant *Fam92a* (Fig. 4a), indicating that the phenotype is both specific and independent of the blasticidin-based readout. Notably, ciliogenesis was also severely reduced in *Fam92a* knockout cell pools (Fig. 4b). To gain further insight into *Fam92a* function, we identified FAM92A-associated proteins using a cell line expressing FAM92A-LAP (localization and affinity purification tag consisting of S-tag-HRV3C-GFP). Affinity purification of FAM92-LAP specifically recovered the known transition zone components CBY1 and DZIP1L (Fig. 4c and Supplementary Table 6)^35-37^. Consistent with this finding, we observed prominent FAM92A localization at the transition zone using both an antibody whose specificity we validated in knockout cells (Supplementary Fig. 4b-c) and the FAM92A-LAP cell line (Fig. 4d). While this work was in progress, another group independently identified FAM92A as a transition zone protein contributing to ciliogenesis, corroborating our studies^38^.

To characterize the TPR domain-containing protein TTC23, we analyzed TTC23- interacting proteins by affinity purification and mass spectrometry. Notably, the most prominent TTC23-associated proteins were IQCE and EFCAB7, which represent the non-transmembrane components of the Ellis-van Creveld (EvC) zone. The EvC zone is a proximal region of the cilium that is important for Hh signaling but dispensable for cilium assembly^39,40^ (Fig. 4e and Supplementary Table 6). Of the four proteins–EVC, EVC2, IQCE, and EFCAB7– that are known to localize to the EvC zone, *EVC* and *EVC2* are also mutated in the ciliopathy Ellis-van Creveld syndrome^41^. Consistent with the biochemical interactions we identified, TTC23-LAP co-localized with EVC and IQCE at the EvC zone (Fig. 4f and Supplementary Fig. 4d). To assess TTC23 function, we examined *Ttc23* knockout cells, which showed a moderate reduction in inducible blasticidin resistance but no defect in cilium assembly (Supplementary Fig. 4a,e). *Ttc23* knockout cells also showed reduced localization of IQCE and EVC to the EvC zone (Fig. 4g and Supplementary Fig. 4f-g); conversely, *Iqce* RNAi led to a strong decrease in TTC23-LAP localization to the EvC zone (Fig. 4h). Together these results establish TTC23 as a novel EvC zone component that participates in Hh signaling.

**Figure 4.**
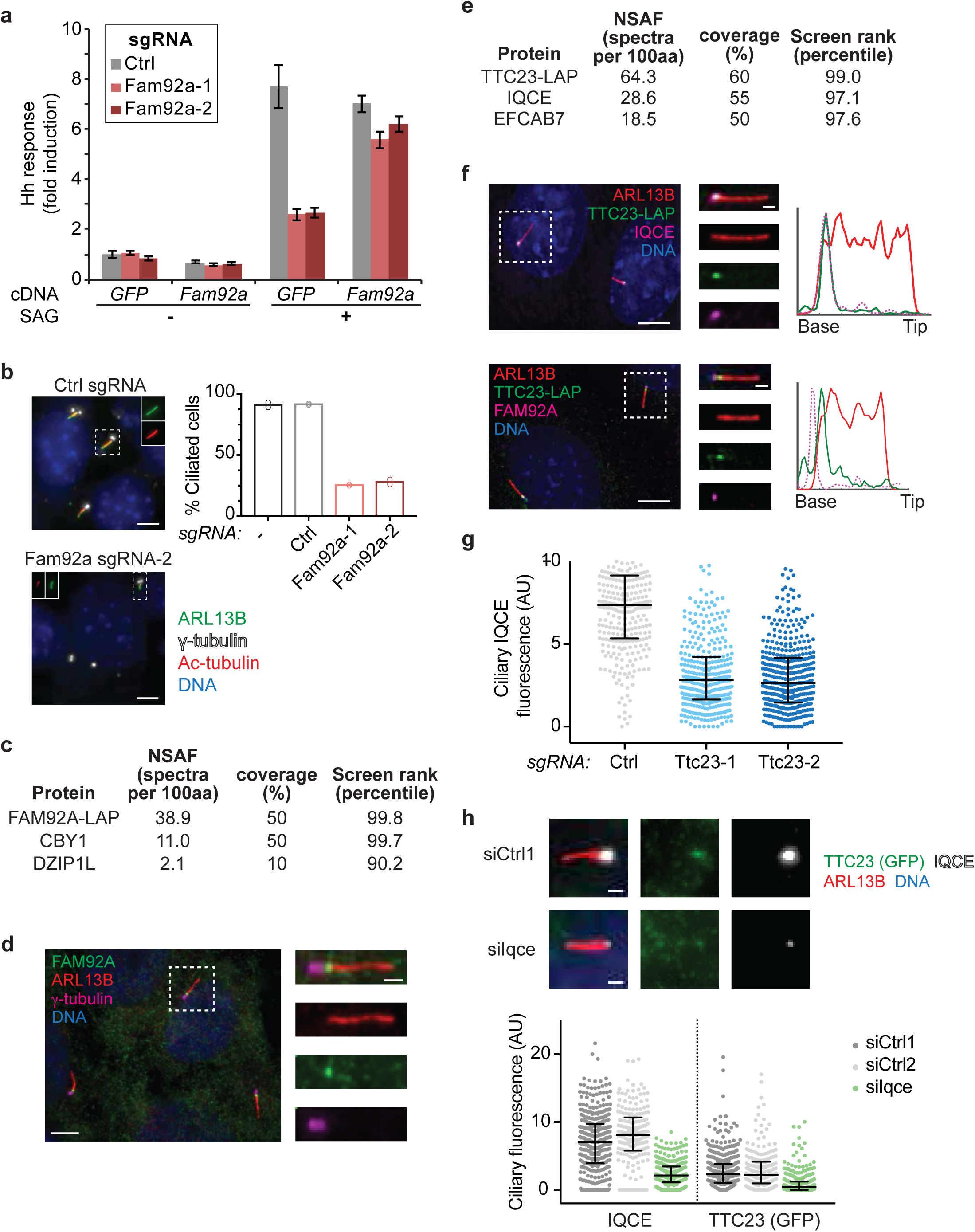
Characterization of FAM92A and TTC23 as transition zone and EvC zone components. **a**) A luciferase-based Hh pathway reporter assay was performed for cells transduced with the indicated sgRNAs and transfected with plasmids expressing either *Fam92a*-3xFLAG (*Fam92a*) or *GFP-FKBP* (*GFP*). Cells were untreated (-SAG) or stimulated with the SMO agonist SAG (+SAG). Mean and standard deviation for 4 replicate measurements from one of two representative experiments. **b**) 3T3-[Shh-BlastR;Cas9] cells transduced with *Fam92a*-targeting sgRNAs are deficient in cilium assembly, as visualized by immunofluorescent staining of ARL13B and acetylated tubulin; centrioles are marked by γ-tubulin. Bars show mean fraction of ciliated cells; dots show ciliated fraction in each independent experiment (>400 cells were analyzed from two independent experiments). Scale bar: 5 μm **c**) Mass spectrometry analysis of FAM92A-associated proteins purified from IMCD3 cells reveals transition zone components CBY1 and DZIP1L. For each protein, the normalized spectral abundance factor (NSAF; number of spectra identified per 100 amino acids), the percent of the protein covered by identified peptides, and the percentile rank of the corresponding gene in the screen dataset are indicated. **d**) FAM92A (green) localizes to the transition zone of IMCD3 cells, distal to centrioles marked by γ-tubulin (magenta) and proximal to the ciliary shaft marked by ARL13B (red). Scale bars: 5 μm and 1 μm (insets). **e**) Mass spectrometry analysis of TTC23-associated proteins purified from IMCD3 cells reveals EvC zone components EFCAB7 and IQCE. Data are shown as in (**c**). **f**) TTC23-LAP stably expressed in IMCD3 cells co-localizes with EvC zone component IQCE, distal to transition zone component FAM92A. Line plots show normalized intensity for the indicated markers along the length of the cilium; tick marks are 1 μm intervals. Scale bars: 5 μm and 1 μm (insets). **g**) Ciliary IQCE levels were measured for cells transduced with control (Ctrl) sgRNA or sgRNAs targeting *Ttc23*. The median and interquartile range are plotted for N > 295 cells measured in each condition from a representative experiment out of two independent replicates. **h**) Knockdown of *Iqce* in IMCD3 cells leads to reduced ciliary GFP (TTC23-LAP) and IQCE (top). The median fluorescence and interquartile range are plotted for n> 190 cilia from a single experiment out of two (IQCE) or four (GFP) replicate experiments (bottom). Scale bars: 1 μm.

### Identification of novel disease genes

Because the vast majority of ciliopathy genes were hits in the screen, we asked whether uncharacterized hit genes may be mutated in ciliopathies of previously unknown etiology. We first examined *Txndc15*, a screen hit that encodes a thioredoxin domain-containing transmembrane protein of unknown function. Notably, a previous analysis of MKS patients identified a family with a *TXNDC15* mutation^42^. However, other candidate genes could not be ruled out, and an *EXOC4* variant was favored as the causative mutation. To investigate whether loss of *TXNDC15* might cause a ciliopathy, we analyzed *Txndc15* knockout cells using the luciferase-based reporter assay, finding a clear defect in Hh signaling. Furthermore, introduction of wildtype *Txndc15* rescued this defect, whereas cDNA encoding the mutant allele found in MKS patients behaved like a null allele (Fig. 5a). We also found that cilia in *Txndc15* knockout cells exhibited increased variability in length and decreased levels of the ciliary GTPase ARL13B (Fig. 5b and Supplementary Fig. 5a-b). Thus, *TXNDC15* likely represents a novel MKS gene, and our screening system can provide a straightforward means to functionally test candidate mutations identified in patients. Consistent with our findings, a very recent follow-up study has identified additional MKS families with *TXNDC15* mutations^43^.

Similarly, the finding that *Armc9* is a hit in our screen raises the possibility that it is a causative ciliopathy gene. Recently, Kar et al. reported that individuals with a homozygous splice site mutation in *ARMC9* present with mental retardation, polydactyly, and ptosis^44^ but rejected a diagnosis of Bardet-Biedl syndrome^45^. We found that cilia from *Armc9^-/-^* cells were short and exhibited reduced levels of acetylated and polyglutamylated tubulin (Fig. 5c and Supplementary Fig. 5c). Furthermore, ARMC9-3xFLAG prominently localized to the proximal region of cilia when stably expressed in IMCD3 cells. Notably, stimulation of Hh signaling led to relocalization of ARMC9 towards the ciliary tip within 6 hr before a gradual return to its original proximally biased localization, suggesting that ARMC9 may become ectocytosed at later time points^46^ (Fig. 5d-e). Furthermore, signaling-induced tip accumulation of GLI2 and GLI3 was reduced in *Armc9* mutant cells (Fig. 5f and Supplementary Fig. 5d) but SMO translocation to cilia was intact (Supplementary Fig. 5e), suggesting that ARMC9 specifically participates in the trafficking and/or retention of GLI proteins at the ciliary tip. Collectively, these findings demonstrate that ARMC9 is a ciliary signaling factor and suggest that *ARMC9* is a novel ciliopathy gene.

**Figure 5.**
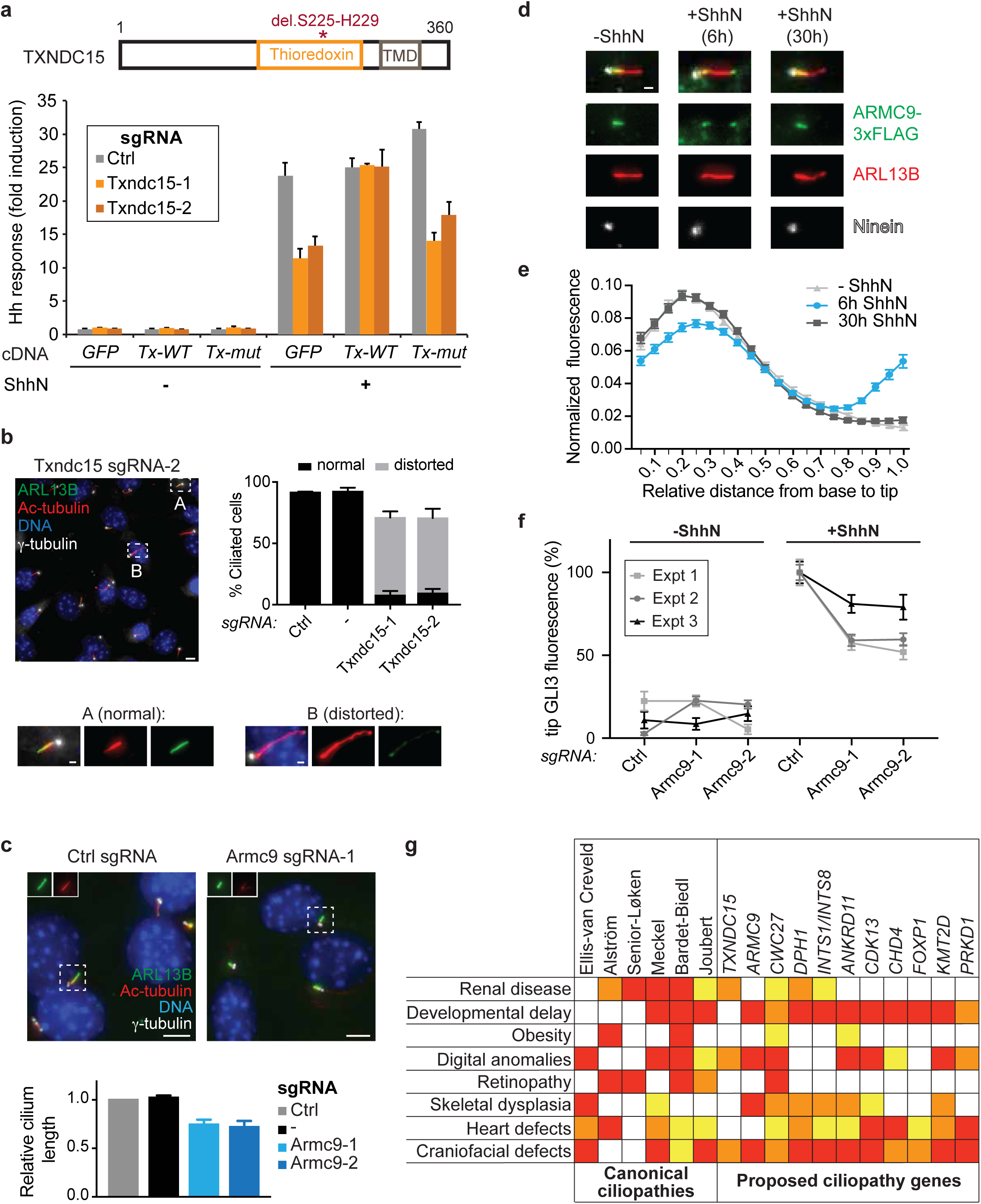
Insights into ciliopathies from previously uncharacterized screen hits. **a**) Diagram of TXNDC15, with the transmembrane domain (TMD), thioredoxin domain, and 5 amino acid deletion found in MKS patients indicated (top). Hh luciferase reporter assay for cells transduced with the indicated sgRNAs and transfected with plasmids encoding either *GFP-FKBP(GFP)*, wildtype *Txndc15* (*Tx-WT*), or mutant *Txndc15* (*Tx-mut*). Cells were untreated or stimulated with ShhN. Mean and standard error for 6 replicate measurements from one of two representative experiments. **b**) Analysis of cilia in 3T3-[Shh-BlastR;Cas9] cells transduced with *Txndc15*-targeting sgRNAs reveals many distorted cilia with abnormal morphology or length and decreased ARL13B staining. Lower images show enlarged micrographs for cilia marked by boxes A and B. Bar graph shows mean and standard deviation (three independent experiments, >700 cells counted total). Scale bars: 5 μm (upper images) and 1 μm (lower images). See also Supplementary Figure 5a-b. **c**) Analysis of cilia in 3T3-[Shh-BlastR;Cas9] cells transduced with *Armc9*-targeting sgRNAs reveals shorter cilia and reduced staining for the ciliary marker acetylated tubulin. Representative micrographs are shown at top; mean cilium length relative to control sgRNA and standard deviation are shown at bottom (three independent experiments, >300 cilia analyzed per experiment). Scale bars: 5 μm. See also Supplementary Figure 5c. **d**) Analysis of ARMC9-FLAG localization along IMCD3 cilia following stimulation of Hh signaling. Representative micrographs showing localization relative to centrioles, marked by ninein, and the cilium, marked by ARL13B. Scale bar: 1 μm. **e**) Plots of ARMC-FLAG intensity along the length of the cilium from base (position 0) to tip (position 1.0) are shown for IMCD3 cells treated as indicated. The mean and standard deviation are plotted after normalizing the total intensity in each cilium to 1.0 (N > 500 cilia, representative example from 3 independent experiments). **f**) Fluorescence intensity of GLI3 at the cilium tip was measured for the indicated cells in the presence or absence of ShhN. Mean intensity (relative to control sgRNA cells +ShhN) and standard error of the mean are shown for each of three experiments (N > 250 cilia in each experiment). See also Supplementary Fig. 5d. **g**) Table showing select clinical features in a diverse set of ciliopathies and their observation in the context of specific mutations and syndromes. Colors indicate high (red), moderate (orange) and low (yellow) prevalence. The name of each syndrome is listed; Joubert encompasses Joubert Syndrome and related disorders.

The observation that *ARMC9-* (and *DPH1-*) based syndromes likely represent unrecognized ciliopathies led us to ask whether our screen could help classify other genetic disorders as ciliopathies. Consistent with this possibility, the peptidyl-prolyl isomerase *Cwc27* was a hit in our screen and was recently identified as a retinitis pigmentosa (RP) gene^47^. Although this syndrome was not recognized as a ciliopathy, the spectrum of *CWC27* pathologies reported includes canonical ciliopathy symptoms (craniofacial abnormalities, short stature, brachydactyly, and developmental delay), and we therefore suggest that *CWC27*-associated disease is a ciliopathy. Similarly, mutations in genes encoding the *INTS1* and *INTS8* subunits of the Integrator complex were recently described in individuals with a neurodevelopmental disorder with additional facial and skeletal malformations commonly seen in ciliopathies^48^. As the Integrator genes *Ints6* and *Ints10* are hits in our screen, disorders due to defects in Integrator complex function may also stem from altered ciliary signaling.

These cases of individual disorders that can now be classified as likely ciliopathies led us to ask whether systematic efforts to map disease genes might reveal broader commonalities with our screen hits. Strikingly, we found that screen hits *ANKRD11*, *CDK13*, *CHD4*, *FOXP1*, *KMT2D*, and *PRKD1* were all recently identified in a large-scale exome sequencing study of patients with congenital heart defects (CHD)^49^. The significant overlap in these two unbiased datasets (*P* = 6.11 x 10^-4^) provides compelling evidence that defective ciliary signaling may be a prevalent cause of CHDs. Moreover, mutations in these genes appear to cause *bona fide* ciliopathies, as patients also exhibit ciliopathy symptoms including craniofacial abnormalities and developmental delay (*ANKRD11*, *CDK13*, *CHD4*, *FOXP1*, *KMT2D*, *PRKD1*), dysgenesis of the corpus callosum (*CDK13*), polydactyly (*CHD4*), obesity (*ANKRD11*), and craniofacial malformations (*ANKRD11* and *KMT2D*)^4,49,50^ (Fig. 5g). A link between primary cilia and heart defects was reported in a recent forward genetic study in mice^51^, and our screen now extends this finding to include several human disease cases.

### A new protein complex for centriole stability

Unexpectedly, our screen hits encode not only canonical ciliary proteins but also centriolar proteins not previously linked to ciliary Hh signaling. Given the roles of hits such as CEP19, CEP44, CEP120, and CEP295 at centrioles, we considered whether other hits might also have centriolar functions. Indeed, the uncharacterized hit 1600002H07Rik (human C16orf59) localized to centrioles when stably expressed in IMCD3 cells (Fig. 6a). We performed affinity purifications and found that 1600002H07Rik-LAP co-purified with the uncharacterized protein 4930427A07Rik (human C14orf80) and the distant α/(β-tubulin relatives ε-tubulin (TUBE1) and δ-tubulin (TUBD1) (Fig. 6b). All four of these gene products are also among the top-scoring hits in our screen, and ε-tubulin and δ-tubulin have previously been linked to centriole assembly and maintenance^52-56^. We therefore propose to name *4930427A07Rik*/*C14orf80* and *1600002H07Rik*/*C16orf59* as *Ted1* and *Ted2*, respectively, for their functional link and biochemical association with tubulins epsilon and delta.

**Figure 6.**
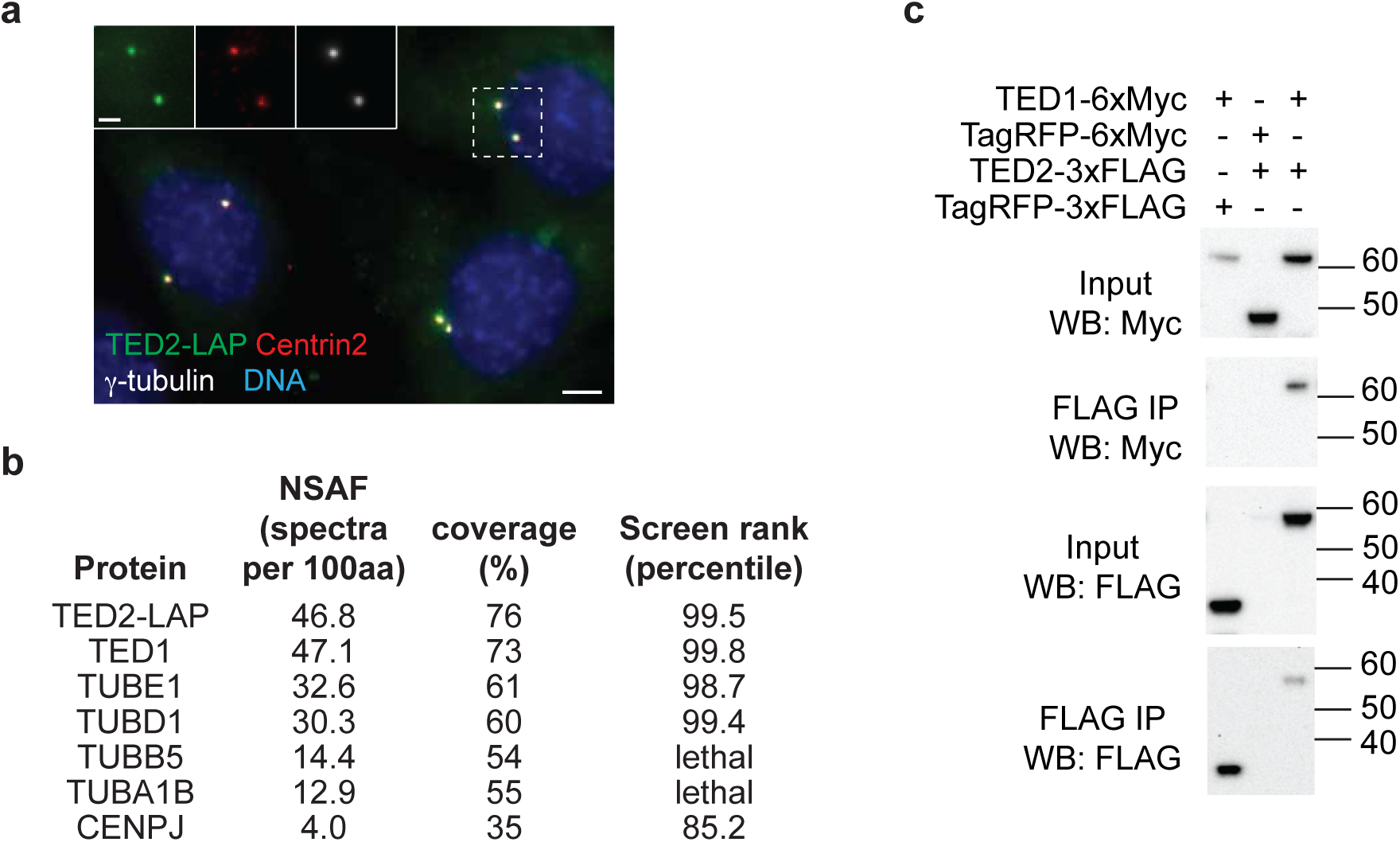
TED1 and TED2 form a tetrameric complex with δ- and ε-tubulins. **a**) IMCD3 cells stably expressing TED2-LAP were immunostained with antibodies to centrin2 and γ-tubulin to visualize centrioles. Scale bar: 5 μm (2 μm for insets). **b**) Mass spectrometry analysis of TED2-associated proteins purified from IMCD3 cells reveals TED1, ε-tubulin, δ-tubulin in nearly stoichiometric amounts, as well as α/(β-tubulin and CENPJ. For each protein, the normalized spectral abundance factor (NSAF; number of spectra identified per 100 amino acids), the percent of the protein covered by identified peptides, and the percentile rank of the corresponding gene in the screen dataset are indicated. **c**) Binding of TED1 and TED2 was assessed via co-immunoprecipitations performed in HEK293T cells transfected with the indicated proteins. Recovered proteins were analyzed by Western blot.

Our mass spectrometry analysis of TED2-associated proteins revealed approximately stoichiometric amounts of TED1, TED2, ε-tubulin, and δ-tubulin, as seen by comparison of the normalized spectral abundance factors (Fig. 6b and Supplementary Table 6). We also noted lower amounts of co-purifying α/β-tubulin and CENPJ/CPAP, a centriolar regulator of microtubule dynamics^57-59^. To corroborate these mass spectrometry analyses, we confirmed co-purification of TUBD1 and TUBE1 by Western blot, readily detecting these proteins in our TED2-LAP purification but not in a negative control purification (Supplementary Fig. 6a). We also tested for an interaction between TED1 and TED2 by co-transfection and co-immunoprecipitation. Not only did we detect a specific interaction between these proteins, we found that TED1 and TED2 mutually stabilize their expression levels, leading to notably higher expression when expressed together versus individually (Fig. 6c).

To functionally characterize *Ted1* and *Ted2*, we first examined knockout cells generated using two sgRNAs targeting each gene. Strikingly, we found that *Ted1* and *Ted2* knockout cell pools were almost completely devoid of centrioles, as assessed by staining with antibodies to centrin, ninein, polyglutamylated tubulin or γ-tubulin (Fig. 7a-b). Congruently, mutant cells also lacked cilia (Fig. 7a) and had strong defects in Hh pathway reporter induction and GLI protein regulation (Supplementary Fig. 6b-c). We also noted that *Ted1* and *Ted2* mutants exhibited a mild growth defect (Supplementary Table 3), which is consistent with recent evidence that NIH-3T3 cells lacking centrioles can continue to proliferate, albeit at a reduced rate^60^. By contrast, in other cell types, a p53-dependent arrest prevents proliferation in the absence of centrioles. These observations prompted us to investigate whether loss of *Ted1* or *Ted2* has varying effects on proliferation across different cell types, and whether such phenotypic variability could enable predictive identification of genes with similar function. We therefore examined a recently published collection of CRISPR-based growth screens conducted in 33 different human cell lines and used hierarchical clustering to group genes based on the similarity of their cell type-specific growth phenotypes^61^. Strikingly, this unbiased approach placed *Ted1*, *Ted2*, and *Tube1* in a single cluster (Fig. 7b and Supplementary Table 7), suggesting that they have a highly similar function in addition to encoding components of a shared protein complex. We note that *TUBD1* did not co-cluster but was likely not effectively targeted by this sgRNA library.

**Figure 7.**
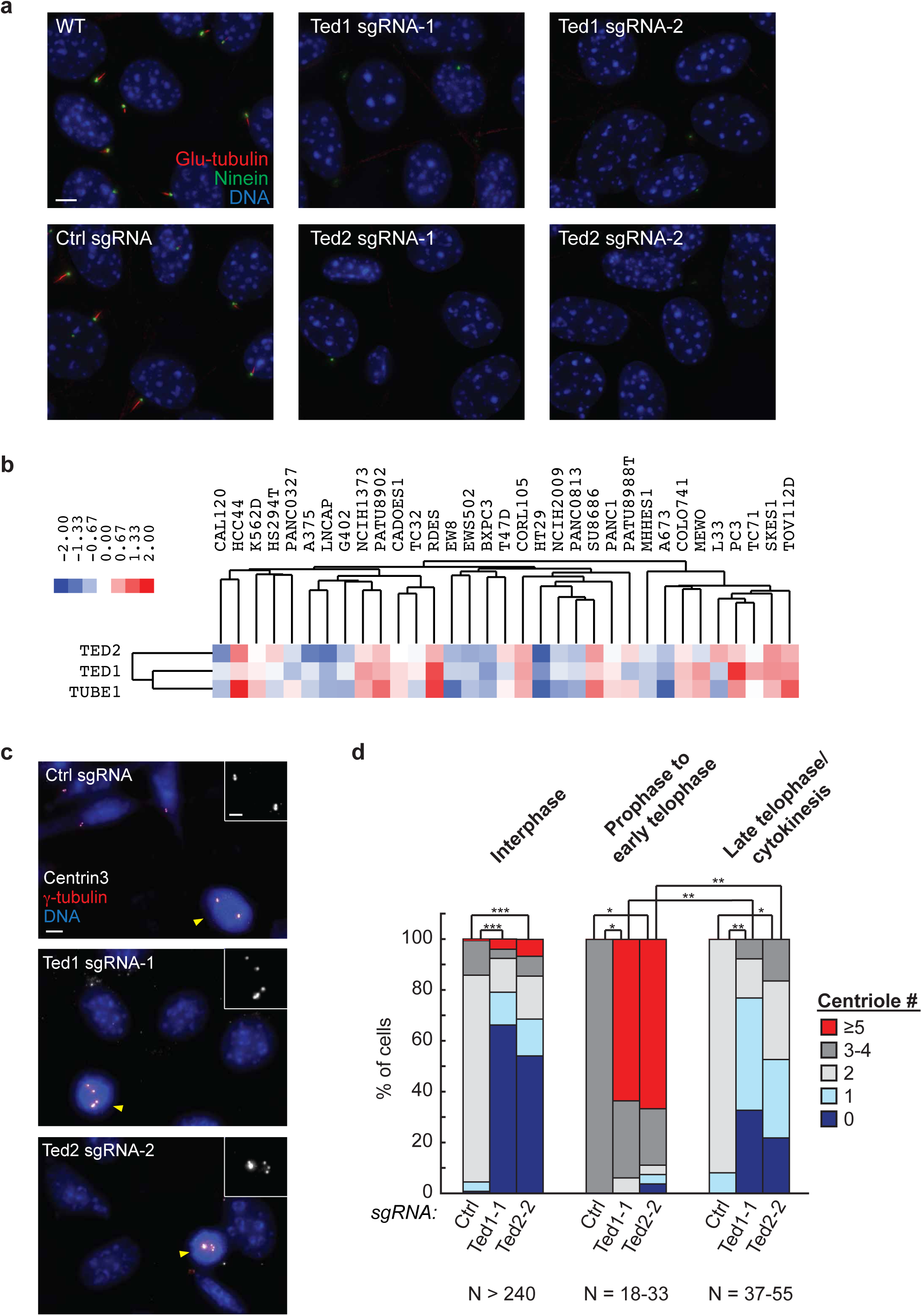
The TED complex is required for centriole stability. **a**) Cells transduced with the indicated sgRNAs were stained with antibodies to ninein (centrioles) and polyglutamylated tubulin (GT335, centrioles and cilia). Scale bar: 5 μm. **b**) Hierarchical clustering of relative growth scores across the indicated cell lines reveals that *TED1*, *TED2*, and *TUBE1* share a similar pattern of relative fitness. Blue and red shading indicates decreased and increased proliferation relative to the average behavior across all cell lines. **c**) For cells transduced with the indicated sgRNAs, centrioles were visualized by staining with antibodies to centrin3 and γ-tubulin. Insets show centrin3 staining in mitotic cells, marked by yellow arrowheads. Scale bars: 5 μm (2 μm for insets). See also Supplementary Figure 6d. **d** Centrioles marked by centrin3 and γ-tubulin were counted in cells at the indicated cell cycle stages, with each cell categorized as having either zero centrioles, some centrioles but fewer than expected for that stage in the cell cycle (two or four), the expected two or four centrioles, or more than the expected number of centrioles. Statistically significant differences in centriole counts are shown for select conditions (*, *P* < 1x10^-6^; **, *P* < 1x10^-10^; ***, *P* < 1x10^-60^, determined by Fisher’s exact test; see **Methods**).

To better understand the basis for centriole loss in *Ted1* and *Ted2* mutants, we examined centrioles in cells at different stages of the cell cycle. Surprisingly, while mutant cells typically had zero or one centriole in interphase, nearly all mitotic cells had excess centrioles (more than four), suggestive of *de novo* centriole formation before mitotic entry (Fig. 7c-d and Supplementary Fig. 6d). By contrast, cells exiting mitosis more closely resembled interphase cells, again showing significantly decreased centriole numbers compared to negative control sgRNA-expressing cells (Fig. 7d). Taken together, these observations suggest that *Ted1* and *Ted2* are dispensable for centriole biogenesis but required for centriole stability, with newly generated centrioles rapidly lost as cells exit mitosis.

While the biochemical activities of ε-tubulin and δ-tubulin remain elusive, very recently *TUBE1*- and *TUBD1*-deficient human RPE1 cells were found to frequently lack centrioles and undergo cycles of *de novo* centriole biogenesis followed by disappearance of these centriole-like structures during progression through the cell cycle (J. Wang and T. Stearns, personal communication). This phenotype echoes our findings in *Ted1* and *Ted2* mutant cells, and together with our biochemical findings indicates that the TED complex is required for centriole stability.

## Discussion

Here we present a new functional screening platform that pairs a pathway-specific selectable reporter with genome-wide CRISPR-based gene disruption. Applying these technologies to systematically investigate cilium-dependent Hh signaling, we obtain a comprehensive portrait of cilium biology that identifies hit genes with high sensitivity and specificity, surpassing the performance seen in related siRNA-based arrayed screens. As a result, we identify many novel hit genes, and we show that a number of previously uncharacterized gene products have diverse functions in cilium assembly and ciliary signaling.

### Screen performance and CRISPR-based screening

Several factors likely contributed to the quality of our screen, including the pooled screening format, the use of CRISPR for gene disruption, and a newly designed sgRNA library. Given the strong influence of cell confluence on cilium assembly and Hh signaling, the homogeneous growth conditions afforded by pooled screening were likely a major advantage. In an arrayed format, perturbations causing growth defects can indirectly affect ciliogenesis by decreasing cell density, thus generating false positives that need to be filtered out^10^. Pooled screens achieve confluence regardless of genotype, and thus we can successfully identify hits that have moderate proliferation defects, including the four components of the TED complex.

Another key feature of our screen is the use of CRISPR-based gene disruption. The strong phenotypes produced by CRISPR/Cas9 likely made it possible to detect hit genes in cases where partial knockdown by RNAi or CRISPR interference (CRISPRi) might have failed to produce a detectable phenotype. A potential caveat of our approach is a decreased ability to detect hits among genes that are strictly required for cell viability. However, because ciliary signaling is dispensable for growth of cultured cells, this issue likely had a limited impact on our screen. It is also important to note that the allelic series achievable with RNAi and CRISPRi may be preferable when screening for phenotypic modifiers, as in chemical genetic or genetic interaction analyses.

The strong performance of CRISPR-based screening may also be attributable to our use of an sgRNA library comprised of many highly active sgRNAs with few off-target effects. High on-target activity is especially important for detecting hits in dropout-based screens (in which hits become depleted) and was achieved by using 10 sgRNAs per gene and by optimizing the stability of Cas9 expression. A second benefit of using 10 sgRNAs per gene is the increased statistical power achieved when multiple effective sgRNAs are found targeting a single gene. Indeed, for hit genes such as *Dync2h1*, *Tmem107*, *Ift80*, *B9d1*, and *Grk2*, at least 7 out of 10 sgRNAs were depleted more than 4-fold (and up to 45-fold), leading to high statistical confidence. While other genes may not have been targeted as efficiently, the use of 10 elements per gene strongly increases the statistical power of hit gene detection.

A final important feature of our screening platform is that it can readily be extended to other biological processes. To date, most CRISPR-based screens have been limited to situations where the phenotype of interest directly affects cell viability, such as defining factors affecting cell proliferation or resistance to toxins, drugs, or microorganisms. By using a reporter-based selectable marker, our approach allows powerful screening tools to be applied to virtually any process with a well-defined transcriptional response.

### New insights into cilium biology, Hh signaling, and ciliopathies

The combined benefits of our pooled CRISPR-based screening approach enabled us to generate a rich dataset that we anticipate will be a lasting resource in the field, both for dissecting the basic mechanisms of ciliary signaling and also for defining new ciliopathy genes and potential therapeutic targets in Hh-driven cancers. While prior siRNA-based screens have contributed to our understanding of cilium function and Hh signaling, many of these datasets suffer from false positives or false negatives that limit their utility. In addition, most screens focused on cilium assembly rather than Hh signaling and thus do not identify Hh pathway components. Roosing et al.^9^ improved their ciliogenesis screen results by using known positive and negative gene sets to help classify hits and by incorporating gene expression datasets in addition to their microscopy-based screen data. Our screening approach achieved high performance without dependence on other data sources, which may not always be available, or on *a priori* definition of hits, which could bias discovery of new hit classes. Our screens are also highly reproducible, thereby enabling comparative screening approaches that will be instrumental in uncovering novel factors acting at specific steps in Hh signaling, in particular the regulation of SMO by PTCH1, a poorly understood yet critical aspect of Hh signaling.

The practical value of our screen is demonstrated by the discovery of new genes that participate in ciliary signaling and new candidate ciliopathy genes. While the precise roles of FAM92A at the transition zone and TTC23 at the EvC zone will require further study, our screen demonstrates that new components remain to be identified even for these well-studied ciliary structures. Similarly, our analyses of *TXNDC15*, *ARMC9*, *CWC27*, *DPH1*, *INTS6*, *INTS10*, *ANKRD11*, *CDK13*, *CHD4*, *FOXP1*, *PRKD1,* and *KTM2D* illustrate that screen hits can help to identify a ciliopathy-causing gene from a short list of variants, as is frequently the case in modern genome sequencing of small pedigrees, and to classify new genetic syndromes as disorders of ciliary signaling. Indeed, with the exception of *TXNDC15*, all of the aforementioned genes had been previously linked to disease states (e.g. *KMT2D* and Kabuki Syndrome, OMIM 147920; *ANKRD11* and KBG Syndrome, OMIM 148050; *DPH1* and Loucks-Innes Syndrome, OMIM 616901) without a potential role for cilia described. Among the syndromes caused by mutations in these genes, it is striking that the most prevalent feature is CHD. Our screen thus provides unbiased evidence that several CHD cases are *bona fide* ciliopathies, thereby building upon similar connections observed in mice and motivating future investigations by human geneticists and developmental biologists.

It is also noteworthy that our screen hits include the vast majority of known ciliopathy genes but not those primarily linked to kidney pathology (with or without laterality defects). In particular, the NPHP genes that are associated with the Inversin compartment (*INVS*, *NPHP3*, *ANKS6*, *NEK8*) and the polycystic kidney disease genes *PKD1*, *PKD2*, and *PKHD1* are all non-hits. This finding suggests that these renal diseases are mechanistically distinct from other ciliopathies, likely acting independently of Hh signaling and possibly even involving pathomechanisms independent of cilia^62^. By capturing a comprehensive and unbiased picture of cilium-based signaling, our screen refines the classification of ciliopathies. More broadly, as genome sequencing reveals disease-associated variants at ever-growing rates, genome-wide functional studies such as that presented here will become a powerful resource to distinguish disease-causing mutations from innocuous variants^63^ and to gain insight into underlying disease mechanisms.

Our screen also identifies hit genes with highly diverse roles in cilium function and Hh signaling, including several genes that participate in centriole biology. For hits *Ted1*, *Ted2*, ε-tubulin and δ-tubulin, we found that these four genes act in concert to ensure centriole stability, as evidenced by the association of their gene products in a complex. This complex appears to be a stoichiometric tetramer and demonstrates that ε- and δ-tubulin directly associate. While the structural basis for this interaction awaits further study, it is tempting to speculate that ε- and δ-tubulin may form a heterodimer analogous to α/β-tubulin. In addition to the physical association of TED complex components, deficiency for these genes leads to a remarkably similar pattern of growth phenotypes across cell lines. This observation provides further evidence for a shared function and complements recent work demonstrating the precise functional predictions made possible by CRISPR-based growth profiling^19^.

Similar to the aforementioned pattern of cell-type-specific growth phenotypes, Turk et al.^64^ recently observed a phylogenetic pattern in which the presence or absence of ε-tubulin in a given species predicts whether δ- or ζ-tubulin is also present. The evolutionary co-occurrence of these centriolar tubulins prompted us to ask whether *Ted1* and *Ted2* also share a similar phylogenetic distribution. We found *Ted1* and *Ted2* homologs in annelids and sea urchin and *Ted1* homologs in evolutionarily distant species such as *Paramecium tetraurelia* and *Tetrahymena thermophila* (Supplementary Fig. 7a). Consistent with a conserved functional relationship among TED complex components, all of these species also have ε-, δ- and/or ζ- tubulins; conversely, we did not detect *Ted1* or *Ted2* homologs in any species lacking ε-, δ- and ζ- tubulins. The scope of this analysis was limited by the more rapid divergence seen for *Ted1* and *Ted2* sequences than for ε-, δ- and ζ-tubulins (Supplementary Fig. 7b), but this phylogenetic distribution further supports a shared function and echoes what is seen for γ-tubulin and subunits of the γ-tubulin ring complex.

Notably, studies across diverse species have implicated ε- and δ-tubulins in stabilizing centriolar microtubules. For example, defects in Paramecium or Tetrahymena ε-tubulin or in *Chlamydomonas reinhardtii* δ-tubulin all lead to centrioles lacking the C and/or B tubules of the centriolar triplet microtubules^53,55,56^. Consistent with these reports, we find that *Ted1* and *Ted2* knockout cells are markedly deficient in centriole maintenance. We speculate that the centriolar structures formed in *Ted1/2*-deficient cells are structurally unstable and degenerate during mitotic exit at the time when pericentriolar material (PCM) is stripped from the centrosome and duplicated pairs of centrioles become disengaged. Since removal of the PCM has been recently shown to cause centriolar instability in fly spermatocytes^65^, we consider post-mitotic PCM removal may be a key event contributing to centriole loss in *Ted* mutants. CENPJ may act together with the TED complex to ensure centriole stability, as CENPJ was found in our TED2 purifications and a mutation in *Cenpj/Sas-4* was recently shown to lead to loss of triplet microtubules in *Drosophila* spermatocytes^66^. Lastly, given that several centriolar genes including *CENPJ* cause microcephaly when mutated^67,68^, TED complex components are potential candidate genes for this neurodevelopmental disorder.

In summary, we have developed a functional screening platform that provides a valuable resource for the investigation of long-standing questions in Hh signaling and in the biology of primary cilia. By applying and extending these tools, it may now be possible to systematically define vulnerabilities in Hh pathway-driven cancers, to search for suppressors of ciliopathies that may inform treatment, and to identify modifiers of Hh pathway-inhibiting chemotherapeutics. Integrating this functional genomic approach with complementary insights from proteomics and human genetics promises a rich toolkit for understanding ciliary signaling in health and disease. More broadly, the approach described here makes CRISPR-based screening possible for essentially any biological process with a defined transcriptional output.

## Acknowledgments

We acknowledge members of the Chen and Nachury labs for advice and technical support; S. van Dorp for assistance with image quantification; P. Beachy for NIH-3T3 cells, 8xGli-luciferase reporter plasmid, and ShhN-producing HEK293T cells; K.C. Garcia and G. Crabtree for use of plate readers; C. Bustamante for use of an Illumina sequencer; R. Rohatgi for antibodies to Evc and Iqce; J. Wang and T. Stearns for sharing unpublished results and cDNA for Cby1; and M. Scott for helpful discussions. This project was supported by NIH Pathway to Independence Award K99HD082280 (D.K.B), Damon Runyon Dale F. Frey Award DFS-11-14 (D.K.B.), seed grants from the Stanford Center for Systems Biology (D.K.B., S.H. and G.T.H.) and Stanford ChEM-H (M.C.B.), an NWO Rubicon Postdoctoral Fellowship (S.H.), National Science Foundation Graduate Research Fellowship DGE-114747 (D.W.M.), a Walter V. and Idun Berry Award (K.H.), T32HG000044 (G.T.H.), DP2 HD08406901 (M.C.B.), R01 GM113100 J.K.C.), and R01 GM089933 (M.V.N.). Cell sorting/flow cytometry was done on instruments in the Stanford Shared FACS Facility, including an instrument supported by NIH shared instrument grant S10RR025518-01. Mass spectrometry analyses were conducted in the Vincent Coates Foundation Mass Spectrometry Laboratory, Stanford University Mass Spectrometry and the Stanford Cancer Institute Proteomics/Mass Spectrometry Shared Resource; these centers are supported by Award S10RR027415 from the National Center for Research Resources and NIH P30 CA124435, respectively. We thank Carsten Carstens, Ben Borgo, Peter Sheffield, and Laurakay Bruhn of Agilent Technologies for cilia-focused oligonucleotide sub-libraries.

## Author Contributions

D.K.B., S.H., J.K.C., and M.V.N. conceived the project with advice from M.C.B. D.K.B. and S.H. developed the Hh pathway reporter screening strategy with assistance from B.K.V. D.W.M., K.H., A. L., G.T.H, and M.C.B. provided functional genomics expertise, the genome-wide sgRNA library, and software for screen data analysis. D.K.B. conducted the genome-wide screen and screen data analysis with assistance from S.H. and A.R.K. D.K.B., S.H., and A.R.K. functionally characterized hit genes of interest, analyzed data, and prepared figures. D.K.B., S.H., and M.V.N. wrote the manuscript with assistance from M.C.B. and J.K.C. D.K.B, S.H., G.T.H., M.C.B, J.K.C. and M.V.N. provided funding for the project.

## METHODS

### CONTACT FOR REAGENT AND RESOURCE SHARING

Further information and requests for resources and reagents should be directed to and will be fulfilled by David Breslow.

## EXPERIMENTAL MODELS AND SUBJECT DETAILS

### Cell lines and cell culture

NIH-3T3 and HEK293T cells were grown in high glucose DMEM supplemented with 10% fetal bovine serum (FBS), 2 mM glutamine, 1 mM sodium pyruvate, 10 U/ml penicillin and 10 μg/ml streptomycin. Light-II NIH-3T3 cells^25^ were grown in the same medium except with 10% bovine calf serum (BCS), and IMCD3 cells were grown in DMEM/F12 medium with the same FBS, glutamine, penicillin and streptomycin additives. Serum starvation was done with the same media but with 0.5% FBS for NIH-3T3 cells, 0.5% BCS for Light-II NIH-3T3 cells, and 0.2% FBS for IMCD3 cells. Cells were passaged using 0.05% trypsin-EDTA. IMCD3 FlpIn cells were provided by Peter Jackson. NIH-3T3 cells were obtained from ATCC. HEK239T-EcR-ShhN cells were provided by Philip Beachy. Cells were confirmed to be mycoplasma free by regular testing with the MycoAlert system (Lonza).

## METHOD DETAILS

### DNA cloning

Individual sgRNAs were cloned by ligating annealed oligonucleotides into pMCB306 or pMCB320 digested with BstXI and Bpu1102I (Fermentas FastDigest enzymes, Thermo Fisher). Ligated products were transformed into Mach1-T 1 competent cells (Thermo Fisher) and recovered plasmids were verified by sequencing.

Cilia-focused sgRNA libraries were cloned from oligonucleotide pools (Agilent) as described^26^. Briefly, oligonucleotides were amplified using primer sequences common to each sub-library, digested with BstXI and Bpu1102I, and ligated into pMCB320, followed by transformation into Endura competent cells (Lucigen). DNA was isolated from collected bacterial colonies using a Plasmid Plus Giga kit (Qiagen).

Individual cDNAs encoding genes of interest were amplified from mouse cDNA or commercially available sources (Dharmacon) and cloned into Gateway Entry vectors either via BP clonase-mediated recombination (Thermo Fisher) or using isothermal assembly. Mutations to introduce resistance to sgRNA-mediated cleavage were introduced by isothermal assembly. Plasmids for expression of tagged genes of interest were generated from Entry vectors by LR clonase-mediated recombination (Thermo Fisher) into Destination vectors encoding C-terminal LAP, 3xFLAG, or 6xMyc tags.

Plasmid pHR-Pgk-Cas9-BFP was cloned by digestion of pHR-SFFV-Cas9-BFP (M. Bassik) and replacement of the SFFV promoter with the Pgk promoter amplified from pEFB/FRT-pCrys-^AP^Gpr161^NG3^-NsiI-pPgk-BirA-ER^46^. Plasmid pGL-8xGli-Bsd-T2A-GFP-Hyg was generated in the pGL4.29-[luc2P/CRE/Hygro] vector (Promega) by replacement of the CRE response element with 8xGli binding sites amplified from pGL3-8xGli-Luc (P. Beachy) and of Luc2P with Bsd-T2A-GFP amplified from pEF5B-FRT-DEST-LAP^69^.

### Virus production and cell transduction

VSVG-pseudotyped lentiviral particles were produced by co-transfection of HEK293T with a lentiviral vector (e.g. MCB320) and appropriate packaging plasmids (pMD2.G, pRSV-Rev, pMDLg/RRE for pMCB320 and pMCB306 sgRNAs; pCMV-ΔR-8.91 and pCMV-VSVG for Pgk-Cas9-BFP). Following transfection using polyethyleneimine (linear, MW~25000, Polysciences), virus-containing supernatant was collected 24 h later, filtered through a 0.45 μm cellulose acetate filter, and either used immediately or stored at -80°C. For sgRNA libraries, a second harvest of viral medium was performed 24 h after the initial harvest. For Cas9-containing virus, lentiviral particles were concentrated 20-fold using Lenti-X Concentrator (Clontech) prior to use.

Cells were transduced by addition of viral supernatants diluted to an appropriate titer in growth medium containing 4 μg/ml polybrene (Sigma Aldrich). Following 24 h incubation at 37°C, virus-containing medium was removed; after an additional 24 h, cells were passaged and, where appropriate, selection for transduced cells was commenced by addition of 2.0 μg/ml puromycin (Invivogen). Multiplicity of infection was determined by flow cytometry-based analysis of fluorescent marker expression.

### ShhN production and titering

The HEK239T-EcR-ShhN cell line was used to produce ShhN-containing conditioned medium. Cells were grown to 80% confluence, medium changed to DMEM with 2% FBS, followed by collection of conditioned medium after 48 h and filtration through a Steritop 0.22 μm filter device (EMD Millipore). The titer of ShhN was determined using NIH-3T3 Light-II 8xGli luciferase reporter cells (see *Luciferase reporter assays*), and a concentration approximately two-fold over the minimum dilution needed for full luciferase induction was used for further experiments.

### Flow cytometry and fluorescence-activated cell sorting

Flow cytometry analyses were conducted using a FACSScan (Becton Dickinson) outfitted with lasers provided by Cytek Biosciences. FACS was performed using FACSAria II cell sorters (Becton Dickinson). Flow cytometry and FACS data were analyzed using Flowjo (Treestar) to quantify the intensity of fluorescent protein expression and fraction of marker-positive cells.

### Generation of stable cell lines

The 3T3-[Shh-BlastR;Cas9] reporter cell line was generated by using Lipofectamine 2000 (Thermo Fisher) to transfect NIH-3T3 cells with pGL-8xGli-Bsd-T2A-GFP-Hyg. Following selection for a hygromycin-resistant cell pool, clonal isolates were obtained by limiting dilution cloning and tested for SAG-induced blastidicin resistance. Lentiviral transduction was then used to introduce Pgk-Cas9-BFP, followed by three rounds of sorting the pool of transduced cells for high-level BFP expression. Notably, we found that other promoters either failed to stably express Cas9 after weeks of passaging or produced an excessive level of Cas9 that led to toxicity (data not shown).

Stable cell lines for affinity purification and localization studies were generated using the FlpIn system (Life Technologies) that relies on FRT-mediated recombination into a single FRT site present in IMCD3 FlpIn cells. Plasmids encoding genes of interest in the pEF5B/FRT-DEST-LAP (or 3xFLAG) vector were transfected into IMCD3 FlpIn cells together with pOG44 Flp recombinase (Life Technologies) using X-tremegene 9 (Roche). Recombined cells pools were obtained following selection with blasticidin (Sigma Aldrich) and were used for further analysis.

### Blasticidin reporter assays

3T3-[Shh-BlastR;Cas9] cells were seeded for signaling and grown to confluence. Growth medium was then replaced with serum starvation medium with or without pathway agonist (ShhN conditioned medium at appropriate dilution or SAG (synthesized as described^70,71^) at 250 nM). After 24 h, cells were passaged at approximately 1:6 dilution to fresh medium with 10% FBS, allowed to adhere for 5 h, and then subjected to blasticidin (blasticidin S hydrochloride, Sigma Aldrich) selection at concentrations ranging from 0.5 to 20 μg/ml for 4 d. Relative viability was determined using the CellTiter-Blue fluorescence-based assay (Promega), and fluorescence read on a SpectraMax Paradigm (Molecular Devices) or Infinite M1000 (Tecan) plate reader.

### Genome-wide screening

Genome-wide screening was conducted in four batches of ~45,000-70,000 sgRNAs^26^. For each batch, lentivirus was produced and titered as described above. 3T3-[Shh-BlastR;Cas9] cells were grown in 15 cm plates and transduced at a multiplicity of infection of ~0.3 in sufficient numbers such that there was a ~500:1 ratio of transduced cells to sgRNA library elements. Cells were selected with puromycin for 5 d, grown for 3 d without puromycin, and then plated for signaling, maintaining a ~ 1,000:1 ratio of cells to sgRNAs for these and all subsequent steps. After cells reached confluence, signaling was initiated by addition of serum starvation medium containing ShhN. After 24 h, cells were passaged, allowed to adhere, and then subjected to blasticidin selection for 4 d at 5 μg/ml, a concentration we found sufficient to achieve strong enrichment/depletion of hits without causing sgRNA library bottlenecks due to excess cell death. After passaging cells to blasticidin-free medium, a ‘T1’ sample was harvested (1000-fold more cells than sgRNAs) and remaining cells were passaged once more before seeding for a second round of signaling and selection. The final ‘T2’ cell sample was collected following 4 d blasticidin selection and one additional passage in the absence of blasticidin. Unselected control cells were also propagated through the entire experiment and harvested at equivalent timepoints.

Screens using the cilia/Hh pathway-focused library were conducted as above except that a variant blasticidin reporter cell line was used in which Cas9-BFP was expressed using the shortened EF1α promoter rather than the Pgk promoter. Because some Cas9-negative cells accumulated during the course of the experiment, the final blasticidin-selected and unselected cells were FACS-sorted for BFP-positive cells bearing Cas9.

To process cell samples for sgRNA sequencing, genomic DNA was first isolated using a QiaAmp DNA Blood Maxi kit or QiaAmp DNA mini kit (Qiagen) depending on the scale of the screen batch. Genomic DNA was then amplified using Herculase II polymerase (Agilent) as described^17^, first using outer primers to amplify the sgRNA cassette from nearly all genomic DNA recovered, then inner primers to amplify a portion of the initial PCR product while introducing sample barcodes and adapters for Illumina sequencing (Supplementary Table 9). Gel purified PCR products were quantified by Qubit 2.0 fluorometer (Thermo Fisher) using the dsDNA HS kit (Thermo Fisher) and pooled for sequencing. Deep sequencing was performed on a NextSeq 500 sequencer with high-output v2 kits (Illumina) to obtain an approximately 500-fold excess of reads to sgRNA library elements. Sequencing was performed using custom primers to read the sgRNA protospacer and sample barcode.

### Luciferase reporter assays

Luciferase reporter assays were conducted using 3T3-[Shh-BlastR;Cas9] cells co-transfected with pGL3-8xGli-Firefly-luciferase and pGL3-SV40-Renilla-luciferase. Cells were transfected using Mirus TransIT-2020 (Mirus Bio) 24 h after plating with luciferase plasmids and either control GFP-containing plasmid (pEF5B-FRT-GFP-FKBP)^72^ or a plasmid encoding a gene of interest. Nearly confluent cells were switched to serum starvation medium with or without pathway agonist 24 h later, and allowed to signal for 24-30 h. Alternatively, luciferase assays for titering ShhN-conditioned medium were performed with the NIH-3T3 Light-II cell line, which has stably integrated versions of GLI-driven firefly luciferase and constitutively expressed Renilla luciferase reporters. After signaling, cells were lysed in lysis buffer (12.5 mM Tris pH 7.4, 4% glycerol, 0.5% Triton X-100, 0.5 mg/mL BSA, 1 mM EGTA, 1 mM DTT) and a dual luciferase measurement performed using a Modulus microplate luminometer (Turner Biosystems).

### Immunofluorescence and localization studies

For localization studies, IMCD3 FlpIn or 3T3-[Shh-BlastR;Cas9] cells were first plated on acid-washed 13mm round #1.5 coverslips (additionally coated with poly-L-lysine for NIH-3T3 cells). After 24 h, cells were transfected with siRNAs (see Supplementary Table 9 for sequences) using Lipofectamine RNAiMAX (Thermo Fisher) or with plasmids using Fugene 6 (Promega). Cells were serum starved for 24 h if needed and then fixed using 4% paraformaldehyde, ice-cold methanol, or both in succession. For GLI2/GLI3/SMO trafficking assays, cells were serum-starved for 20 h, followed by 5-6 h incubation in the presence or absence of the appropriate amount of ShhN-conditioned medium. For analysis of TED2-LAP localization, cells were pre-extracted prior to methanol fixation via a one-minute exposure to PHEM buffer (60 mM PIPES, 25 mM HEPES, 4 mM MgSO_4_, 10 mM EGTA, pH 7.0) with 0.2% TritonX-100.

Fixed coverslips were blocked using PBS with 5% BSA and 5% normal donkey serum, permeabilized if needed using PBS with 0.1% Triton X-100, and then incubated with appropriate primary and secondary antibodies (see Supplementary Table 8 for primary antibodies used). Coverslips were then either stained with Hoechst DNA dye, and mounted on slides using Fluoromount-G mounting medium (Electron Microscopy Sciences), or directly mounted using ProLong Gold antifade reagent with DAPI (Life Technologies).

Coverslips were imaged at 60x or 63x magnification using one of three microscope systems: an Axio Imager.M1 (Carl Zeiss) equipped with SlideBook v6 software, an LED light source (Intelligent Imaging Innovations) and a Prime 95b sCMOS camera (Photometrics); an Axio Imager.M1 (Carl Zeiss) equipped with SlideBook v5 software, a Lambda XL light source (Sutter instruments) and CoolSNAP HQ^2^ CCD camera (Photometrics), or a DeltaVision Elite imaging system equipped with SoftWoRx software, an LED light source, and sCMOS camera (Applied Precision). Z-stacks were acquired at 250-500 nm intervals and deconvolved as needed using Slidebook 6.0 or SoftWoRx softwares.

### Co-transfection and co-immunoprecipitation

HEK293T cells were co-transfected with *Ted1*, *Ted2* or *TagRFP* plasmids using Fugene 6, collected after 48 h, and lysed on ice in CoIP buffer (50 mM Tris pH 7.4, 150 mM NaCl, 1% Triton X-100, 1X DTT, 1X LPB) supplemented with protease inhibitors. Lysates were cleared by centrifugation at 20,000 x *g* for 20 min, and FLAG-tagged proteins were captured by incubation for 2 h with anti-FLAG M2 antibody (Sigma Aldrich) and Protein G Sepharose 4 Fast Flow (GE Healthcare). After four washes of resin with CoIP buffer, bound proteins were eluted by incubation at 95°C in lithium dodecyl sulfate-based gel loading buffer.

### Western blotting

For signaling assays, 3T3-[Shh-BlastR;Cas9] cells transduced with gRNAs as indicated were seeded in a 24-well plate and grown until confluency. Growth medium was then replaced with serum starvation medium with or without ShhN conditioned medium at the appropriate dilution. After 24 h, cells were lysed in SDS sample buffer (50 mM Tris HCl pH 6.8, 8% v/v glycerol, 2% w/v SDS, 100 mM DTT, 0.1 mg/mL bromophenol blue), boiled and sonicated. Samples were loaded onto a 4-15% Criterion TGX Stainfree gel (Bio-Rad), and run for 25 min, 300V in Tris/Glycine/SDS buffer (Bio-Rad), before being transferred onto a PVDF membrane using a Transblot Turbo system (Bio-Rad). Membranes were blocked in 1:1 PBS:SeaBlock (Thermo Scientific) for 1 h at room temperature, and subsequently incubated with the indicated primary antibody for 16 h at 4 °C (primary antibodies to GLI1, GLI2, GLI3, SUFU, and importinβ, as describe d in Supplementary Table 8). Membranes were washed, incubated with HRP-conjugated secondary antibody, washed again, developed using Supersignal West Femto Maximum Sensitivity Substrate (Thermo Fisher) and imaged on a ChemiDoc MP (Bio-Rad). Membranes were stripped using Restore Western Blot stripping buffer (Thermo-Fisher) and re-probed as described above.

For analysis of immunoprecipitations, Western blotting was performed as described above, except samples were separated in 4-12% Bis-Tris PAGE gels (Invitrogen) using MOPS running buffer, transferred to PVDF membranes using the Criterion Blotter system (Bio-Rad), developed using ECL or ECL 2 chemiluminescence detection kits (Pierce), and imaged on a Chemidoc Touch system (Bio-Rad).

### Large-scale affinity purification and mass spectrometry

Affinity purifications were conducted as described^69^. Briefly, approximately 500-1000 μl packed cell volume was lysed in LAP purification buffer containing 0.3% NP-40. Lysate was cleared sequentially at 16,000 x *g* and 100,000 x *g* before incubation with anti-GFP antibody coupled to Protein A resin. After protein capture and washes, bound LAP-tagged proteins were eluted by incubation with HRV3C protease. For mass spectrometry analysis ‘A’ (see Supplementary Table 6), eluted proteins were further purified by capture on S-Protein agarose followed by elution at 95°C in lithium dodecyl sulfate-based gel loading buffer.

For protein analysis by mass spectrometry, gel slices containing affinity-purified proteins were washed with 50 mM ammonium bicarbonate, followed by reduction with DTT (5 mM) and alkylation using propionamide (10 mM). Gel slices were further washed with an acetonitrile-ammonium bicarbonate buffer until all stain was removed. 120 ng of Trypsin/LysC (Promega) reconstituted in 0.1% ProteaseMAX (Promega) with 50 mM ammonium bicarbonate was added to each gel band; after 30 min ~20 μL of additional 50 mM ammonium bicarbonate in 0.1% ProteaseMAX was added. Digestion was then allowed to occur overnight at 37 °C. Peptides were extracted from the gels in duplicate followed by drying using a SpeedVac concentrator. Peptide pools were then reconstituted and injected onto a C18 reversed phase analytical column, ~20 cm in length, pulled and packed in-house. The UPLC was a NanoAcquity or M-Class column (Waters), operated at 450 nL/min using a linear gradient from 4% mobile phase B to 45% B. Mobile phase A consisted of 0.2% formic acid, water, Mobile phase B was 0.2% acetic acid, acetonitrile. The mass spectrometer was an Orbitrap Elite or Fusion (Thermo Fisher) set to acquire in a data-dependent fashion selecting and fragmenting the 15 most intense precursor ions in the ion-trap, where the exclusion window was set at 45 seconds and multiple charge states of the same ion were allowed.

## QUANTIFICATION AND STATISTICAL ANALYSIS

### Analysis of CRISPR-based screens

CRISPR-based screens were analyzed as described^27^, processing data from each screen batch separately. Briefly, to determine sgRNA counts in each sample, raw sequencing reads were trimmed to the 3’-most 17 nt of each protospacer and aligned to expected sgRNA sequences. This alignment was carried out with the makeCounts script of the casTLE software package, which uses Bowtie^73^ to perform alignment with zero mismatches tolerated. The analyzeCounts script (v0.7 and v1.0) of the casTLE software available at https://bitbucket.org/dmorgens/castle^27^ was then used to identify genes exhibiting significant enrichment or depletion and to estimate the phenotypic effect size for each gene. This method uses an empirical Bayesian approach to score genes according to the log-likelihood ratio that a gene’s observed changes in sgRNA counts is drawn from a model of gene effect versus the distribution of negative control sgRNAs. An expected negative score distribution is obtained by random permutation of gene-targeting sgRNA fold-change values and used to determine a *P* value for each gene. Note that the use of 100,000 permutations leads to a minimum reported *P* value of 1 x 10^-5^.

Genes targeted by our sgRNA library that lacked an NCBI identifier or that severely affected growth (casTLE effect size ≤ -2.5 and casTLE *P* value < 0.005) were not considered for further analysis but are included in Supplementary Table 3. Negative and positive reference genes were defined for growth and signaling phenotypes based on gene sets previously defined by Hart et al.^28^ and Roosing et al.^9^, respectively (Supplementary Table 3). Precision-recall and ROC curves using these reference genes were computed in Matlab (Mathworks). The hit genes at 10% and 20% false discovery rate cutoffs were defined using the precision-recall threshold values at precision of 0.9 and 0.8, yielding *P* value cutoffs of 0.0163 and 0.0338, respectively. Ciliopathy-associated genes were defined from the OMIM website. Functional category enrichment analysis for hit genes in the 10% FDR category was performed using the DAVID website’s Functional Annotation Chart tool using all mouse genes as the background^29^. A second analysis was also performed using human homologs of the top hits using all human genes as the background. Significance of overlap between the top 15 congenital heart defect genes reported by Sifrim et al.^49^ and *P* values from functional screening was assessed using the Kolmogorov-Smirnov test.

### Quantification of Hh signaling assays

Blasticidin-based inhibition of cell growth was determined by normalizing raw CellTiter-Blue fluorescence such that growth in the absence of blasticidin corresponds to 100% growth while medium alone corresponds to 0% growth. The IC_50_ for blasticidin was determined using Prism 7.0 (Graphpad Software) to fit dose-response data to log(inhibitor) versus response with variable slope. Data shown are average of two to five independent assays conducted in duplicate for each condition and cell line.

Dual-luciferase data were analyzed by first subtracting background signal such that cells without luciferase give readings of zero. Firefly to Renilla (8x-Gli to constitutive) ratios were then calculated and normalized such that unstimulated wildtype cells have a value equal to 1. Data shown are average of replicate samples from a single experiment representative of 2-3 independent repeats.

### Quantification of fluorescence microscopy images

Microscopy images were analyzed using Fiji ImageJ software (National Institutes of Health) and a custom Matlab (Mathworks) script. Local background subtraction was performed on all images before analysis. To determine ciliary frequency, cells were manually scored for the presence or absence of a cilium using ARL13B and acetylated tubulin as ciliary markers. For ciliary length and intensity of ciliary markers analyses, the ARL13B and/or acetylated tubulin channels were used to create a ciliary mask and to measure cilium length. The ciliary mask was then used to identify and measure ciliary signal in the other channels. The γ-tubulin or ninein signal (staining centrioles) was used to orient all cilia from base to tip. Tip fluorescence for GLI2 and GLI3 was defined as the summed fluorescence in the final five pixels of each cilium, regardless of length. For ARMC9-FLAG localization, the ciliary fluorescence of each individual cilium was normalized to 1, and each axoneme divided in 20 equal distance bins. The fraction of FLAG signal within each bin is plotted for N>500 cilia from one out of three representative experiments. Where indicated in the Figure Legend, data is shown relative to values obtained for the control sgRNA using Prism 7 (GraphPad Software).

Differences in cilium length distribution were tested for significance using the Kolmogorov-Smirnov test in Matlab. Line plots of fluorescence intensity along cilia were generated in ImageJ.

Centriole counting measurements were done manually, using the γ-tubulin and centrin3 staining to guide centriole calling. Cell cycle stage was determined using the DNA morphology, and statistical significance was determined using Fisher’s exact test. Cell counts for each of the five centriole number categories were used for statistical comparisons between genotypes. For comparisons between cell cycle stages, cells observed to have 2 or 3-4 centrioles were grouped into one category to account for the expected reduction in centriole number upon completion of mitosis.

### Analysis of mass spectrometry data

MS/MS data were analyzed using both Preview and Byonic v2.10.5 (ProteinMetrics). All data were first analyzed in Preview to provide recalibration criteria if necessary and then reformatted to MGF format before full analysis with Byonic. Data were searched at 12 ppm mass tolerances for precursors, with 0.4 Da fragment mass tolerances assuming up to two missed cleavages and allowing for fully specific and ragged tryptic peptides. The database used was Uniprot for *Mus musculus* downloaded on 10/25/2016. These data were validated at a 1% false discovery rate using typical reverse-decoy techniques ^74^. The resulting identified peptide spectral matches and assigned proteins were then exported for further analysis using MatLab (MathWorks) to provide visualization and statistical characterization.

### Analysis of CRISPR growth screen datasets

Gene-level growth phenotype data^61^ were downloaded from the Achilles website. Hierarchical clustering using uncentered correlation and average linkage settings was performed using Cluster 3.0 software^75^, and clustered data were visualized in Java Treeview^76^.

### Phylogenetic analysis

Homologs for Tubd1 and Tube1 were either previously described ^64^ or identified using protein BLAST. Homologs for Ted1 and Ted2 were identified using iterative searches with PSI-BLAST^77^. To analyze sequence divergence, homolog sequences were first aligned using Clustal Omega^78^. Phylogenetic trees were generated via neighbor joining with distance correction using Simple Phylogeny ^78^ and visualized using Unrooted^79^.

## DATA AND SOFTWARE AVAILABILITY

Data resources including aligned sgRNA sequences, screen sequencing data, casTLE output, positive and negative reference gene sets for growth and ciliary signaling phenotypes, functional categories enriched among hit genes, phylogenetic analysis of TED complex components, and mass spectrometry-based protein identifications can be found in Supplementary Tables 2, 3, 5, and 6. Software used for casTLE analysis can be found at http://bitbucket.org/dmorgens/castle. Scripts for quantification of cilium intensities and length are available upon request.

## List of Supplementary Items

**Supplementary Table 1. List of select sgRNA sequences.**

**Supplementary Table 2. List of sgRNA counts by deep sequencing of screen cell pools.**

**Supplementary Table 3. casTLE output (gene scores) for genome-wide screen.**

**Supplementary Table 4. Summary of screen data for select genes of interest.**

**Supplementary Table 5. List of functional categories enriched among screen hits.**

**Supplementary Table 6. List of proteins identified by mass spectrometry in affinity purifications.**

**Supplementary Table 7. Clustered growth data from Aguirre et al.**

**Supplementary Table 8. List of primary antibodies.**

**Supplementary Table 9. List of oligonucleotides and recombinant DNA.**

